# Premeiotic endoreplication is the mechanism of obligate parthenogenesis in rock lizards of the genus *Darevskia*

**DOI:** 10.1101/2024.02.27.582286

**Authors:** Dmitrij Dedukh, Marie Altmanová, Ruzanna Petrosyan, Marine Arakelyan, Eduard Galoyan, Lukáš Kratochvíl

## Abstract

Among vertebrates, obligate parthenogenesis occurs exclusively in squamate reptiles. Premeiotic endoreplication in a small subset of developing oocytes has been documented as the mechanism of production of unreduced eggs in minutely explored obligate parthenogenetic lineages, namely in teiids and geckos. The situation in the lacertid genus *Darevskia* has been discussed for decades. Certain observations suggested that ploidy level is restored during egg formation through a fusion of egg and polar body nuclei in *D. unisexualis* and *D. armeniaca*. In this study, we re-evaluated the fusion hypothesis by studying diplotene chromosomes in adult females of sexual species *D. raddei nairensis* and obligate parthenogens *D. armeniaca, D. dahli* and *D. unisexualis*. We revealed 19 bivalents in the sexual species and 38 bivalents in the diploid obligate parthenogens, which uncovers premeiotic endoreplication as the mechanism of the production of non-reduced eggs in parthenogenetic females. The earlier contradicting reports can be likely attributed to the difficulty in identifying mispairing of chromosomes in pachytene, and the fact that in parthenogenetic reptiles relying on premeiotic endoreplication only a small subset of developing oocytes undergo genome doubling and overcome the pachytene checkpoint. This study highlights co-option of premeiotic endoreplication for escape from sexual reproduction in all independent hybrid origins of obligate parthenogenesis in vertebrates studied to date.

## Introduction

It has been estimated that approximately 99.9% of vertebrate species reproduce sexually (Avise, 2009). Rare exceptions to this rule can provide us essential information about the cellular, genetic and evolutionary consequences of sexual reproduction and its loss. Despite its rarity, asexual reproduction in vertebrates is surprisingly diverse and relies on different processes. Gynogenesis, also known as sperm-dependent parthenogenesis, and hybridogenesis, the selective transmission of one intact parental genome to the offspring while the other is renewed by mating, occur in teleost fishes and amphibians (Bogart et al., 2009; Lamatsch and Stöck, 2009, Lavanchy and Schwander, 2019, Stöck et al., 2021a). On the other hand, parthenogenesis, the form of asexual reproduction in which embryo growth and development occur without sperm, has only been documented in cartilaginous fishes (sharks and rays) and sauropsids (non-avian reptiles and birds) (Stöck et al., 2021b).

Facultative parthenogenesis, the type of parthenogenesis in which a female individual can reproduce by both sexual and asexual means, is generally thought to be rare in most cartilaginous fishes and sauropsids. The only species up to now where it appears to be very common is a night lizard of the genus *Lepidophyma* (Kratochvíl et al., 2020). Therefore, in most cases it would be more correct to speak of accidental or spontaneous parthenogenesis, but for simplicity, we use the term facultative parthenogenesis here. In all vertebrates studied, facultative parthenogenesis has been associated with partial or even complete loss of heterozygosity in the offspring (Booth et al., 2012, 2014; Booth and Schuett, 2016; Kratochvíl et al., 2020; Ho et al., 2023). Direct cytological evidence for the process of oogenesis in facultatively parthenogenetic vertebrates is lacking. Based on the observations of the genetic variability in the offspring, it is assumed that the ploidy level of the egg is restored after meiosis by a “fusion” of the nuclei of an egg and a polar body (more likely by the suppression of the second meiotic division) (Booth et al., 2012, 2014; Booth and Schuett, 2016; Kratochvíl et al., 2020), or by doubling of the haploid egg genome leading to the total loss of heterozygosity (Ho et al., 2023).

Among vertebrates, all-female lineages with obligate parthenogenesis occur only in squamate reptiles (lizards and snakes). In most cases studied, their origin is associated with hybridization and the eventual subsequent generation of polyploid clones by mating with a male of a sexual lineage, with the xantusiid genus *Lepidophyma* being a notable possible exception (Sinclair et al., 2010; Wynn et al., 1987; Reeder et al., 2002; Adams et al., 2003; Brunes et al., 2019; Grismer et al., 2014; Abdala et al., 2016; Barley et al., 2021). The mechanism of unreduced egg formation has only been studied in five lizard genera with obligate parthenogenetic species. In the teiid genus *Aspidoscelis* (Cuellar, 1971; Lutes et al., 2010; Newton et al., 2016) and three gecko genera *Hemiphyllodactylus, Heteronotia* and *Lepidodactylus* (Dedukh et al., 2022), the ploidy level is maintained by genome doubling prior to meiosis. During the first meiotic division, bivalents are formed between genetically identical sister chromosomes, produced by the first of two premeiotic replication cycles. The pairing of sister chromosomes ensures the maintenance of heterozygosity. Interestingly, during the pachytene stage of meiosis, two populations of cells can be distinguished (Newton et al., 2016; Dedukh et al., 2022). The first, which is much more numerous, enters the meiosis without premeiotic duplication, fails to form proper bivalents and is arrested by the pachytene checkpoint and driven to apoptosis. Only a small minority of oocytes undergo premeiotic duplication and are able to enter further meiotic phases and to form unreduced eggs (Lutes et al., 2010; Newton et al., 2016; Dedukh et al., 2022).

Premeiotic endoreplication was considered as the mechanism of parthenogenetic reproduction also in several earlier studies of the rock lizards of the lacertid genus *Darevskia* distributed in the area between the Lesser Caucasus Mountains and Lake Van (Uzzell, 1970; Darevsky et al. 1985). For example, Uzzell (1970) noted: “It is possible that the second meiotic division is suppressed in *Lacerta armeniaca*, as suggested by Darevsky (1966), but it seems more likely that there is a premeiotic mitosis without cytokinesis.” However, premeiotic genome endoreplication was further declined and the restoration of the diploid chromosome set by a fusion of the egg cell and the polar body nuclei (central fusion) was again suggested on the basis of observations of pachytene and diplotene during meiosis in adult females of the obligate parthenogenetic diploid species *Darevskia unisexualis* and *Darevskia armeniaca* (Kupriyanova, 2009; Kupriyanova et al., 2021; Spangenberg et al., 2020, 2021). Recently, Spangenberg et al. (2024) reported both non-endoreplicated and endoreplicated oocytes in pachytene of *D. armeniaca*. However, later meiotic stages were not examined and these authors raised questions if postmeiotic fusion could be the mechanism of the production of non-reduced eggs (Spangenberg et al., 2024). It was suggested that oocytes with the original ploidy level overcome a prophase I checkpoint that allows them to proceed in meiosis (Spangenberg et al., 2020, 2024).

Here, we present evidence from the observation of chromosomes in diplotene in a sexual and three parthenogenetic species of the genus *Darevskia*, resolving the long-standing controversy over the mechanism of unreduced egg production in these lizards.

## Material and Methods

### Experimental animals

We compared oocytes at the diplotene stage between adult females of the obligate parthenogenetic hybrid species *D. armeniaca, D. unisexualis* and *D. dahli*, and of *D. raddei nairensis*. representing ancestral, sexual reproduction. *D. unisexualis* evolved by a cross between a female of *D. raddei nairensis* with a male of *D. valentini* (Vergun et al., 2020; Tarkhnishvili et al., 2020). *D. dahli* is a hybrid between a female of *D. mixta* and a male of *D. portschinskii* (Yanchukov et al., 2022; Arekelyan et al., 2022). The origin of *D. armeniaca* is less clear. It was reported to be the product of the hybridization of a *D. mixta* female with a *D. valentini* male (Murphy et al., 2000; Freitas et al., 2019). However, based on allele sharing at microsatellite loci, Tarkhnishvili et al. (2020) suggested that *D. armeniaca* could be the result of a backcross between a parthenogenetic female of *D. dahli* and a male of *D. valentini*, which inherited parthenogenesis from its already parthenogenetic mother.

The experimental animals originated from Armenia (for the list of individuals and their localities see Supplementary Table S1). Alive-animal handling procedures were approved by Yerevan State University according to the ethical guidelines, capture permit No. 3/29.7/1043 was issued by the Ministry of Environment of the Republic of Armenia. The experimental procedures were carried out by an accredited person (M.A., CZ 01223) and under the approval of the Ethical Committee of the Faculty of Science, Charles University (UKPRF/28830/2021).

### Mitotic chromosomes analyses

Mitotic chromosomes were prepared by cultivation of leukocytes from fresh peripheral blood and then standard cytogenetic methods were applied: C-banding for visualisation of constitutive heterochromatin and fluorescence *in situ* hybridization (FISH) with probes detecting telomeres and 18S rDNA clusters. Telomeric FISH using a Cy3-labelled PNA probe was performed according to the manufacturer’s instructions (Telomere PNA FISH Kit/Cy3, Dako), all other methods including the cell cultivation followed the detailed protocols available in Supplementary Data of Altmanová et al. (2023). The 18S rDNA probe labelled by biotin (Roche) has been prepared from genomic DNA of the slow worm *Anguis fragilis*, its efficient hybridization on a broad phylogenetic scale was proven in other sauropsids (Poignet et al. 2021, Altmanová et al. 2023). In both FISH analyses and C-banding, the slides were stained and mounted in Vectashield Antifade Medium containing 4’,6-diamidino-2-phenylindole (DAPI; Vector Laboratories).

### Diplotene chromosomes preparation and analyses

Before the dissection, all experimental animals were sacrificed by blunt force trauma to the head and then the gonads were removed. Diplotene chromosome spreads (also known as lampbrush chromosomes) were prepared according to the protocol of Gall et al. (1991) as described by Dedukh et al. (2022). In order to accurately count chromosomes during diplotene and to assess correct bivalent formation, we performed FISH with the telomeric probe under the same conditions as mentioned above for mitotic chromosomes.

### Widefield and fluorescence microscopy

Microphotographs of mitotic chromosomes were captured by an Axio Imager Z2 microscope equipped with a CoolCube 1 b/w digital camera (MetaSystems) using the MetaSystems platform for automatic search, capture and image processing. Meiotic chromosomes after immunofluorescence staining and FISH were analysed using Olympus BX53 equipped with standard fluorescence filter sets and the microphotographs were captured by a CCD camera (DP30W Olympus) using Olympus Acquisition Software. Microphotographs were finally adjusted and arranged in Adobe Photoshop CS6 software.

## Results and discussion

Cytogenetic inspection of mitotic metaphase chromosomes confirmed previous karyological data in the studied species of the genus *Darevskia* (Kupriyanova, 2009). Females of all species possess diploid chromosome number 2n = 38 with three microchromosomes, one of which is the sex chromosome W (Figures 1, S1). Centromeres, the W chromosome and also terminal regions of some other chromosomes are heterochromatic (Supplementary Figure S2). FISH with the telomeric probe labelled only the ends of the chromosomes (contrary to the report of Kupriyanova, 2009, in *D. armeniaca*), with a stronger signal in the terminal region of the W chromosome in *D. r. nairensis* (Supplementary Figure S2). We found clusters of 18S rDNA on two chromosomes in all the studied species. They are placed on a middle-sized pair in *D. armeniaca* and *D. dahli*, on a smaller pair in *D. r. nairensis*, and on one middle-sized and one smaller chromosome in *D. unisexualis* (Supplementary Figure S2). Compared to sexual *D. r. nairensis*, one cluster was always larger, probably dominant, in the parthenogens, which would explain why only one NOR has been previously reported in the parthenogens (Kupriyanova, 2009). Because the general genome organisation is very similar in all studied species, the apparent cytogenetic evidence of hybrid origin was documented only in *D. unisexualis* thanks to different topology of 18S rDNA between the two parental species chromosome sets.

**Figure 1.**
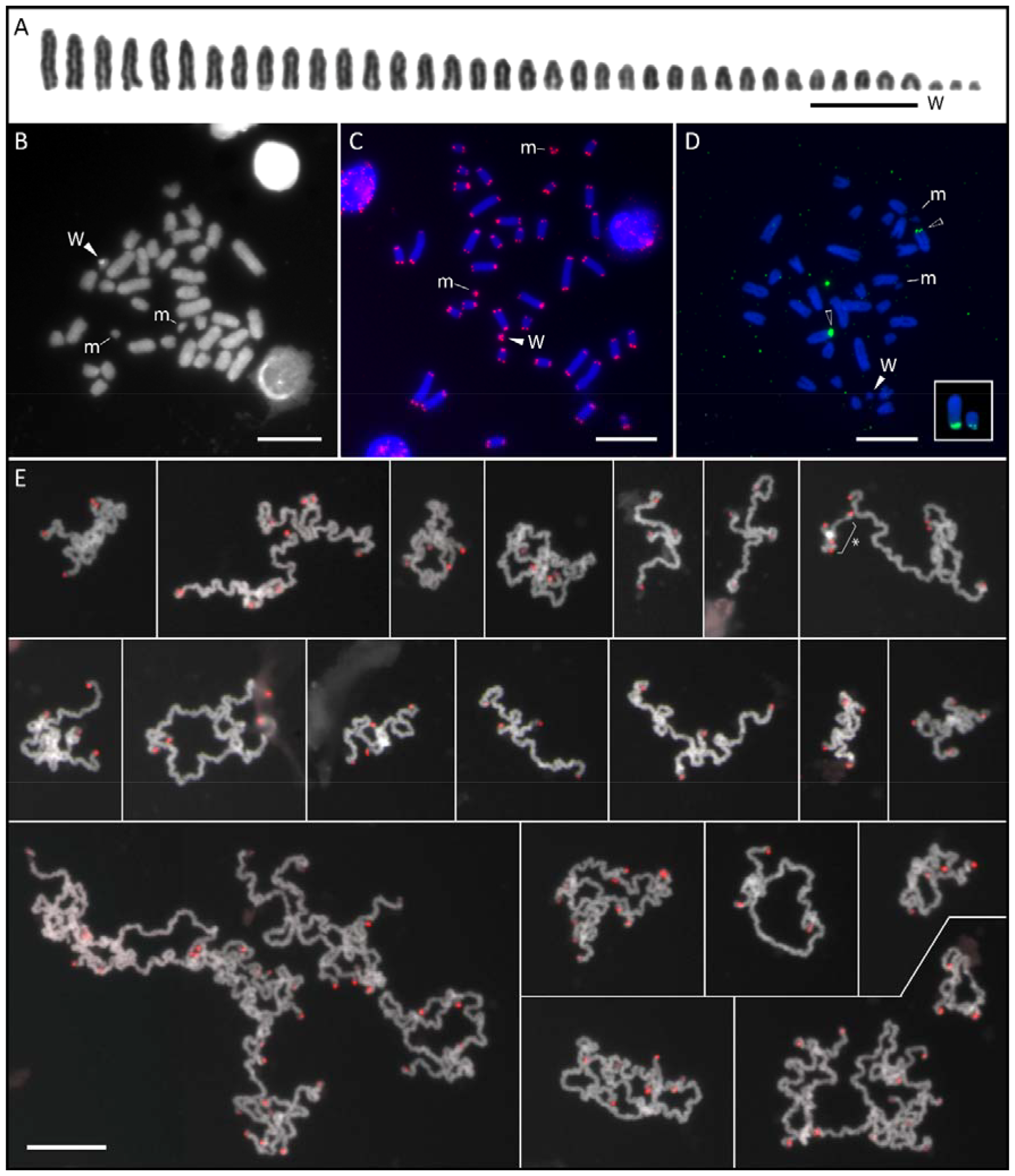
Mitotic and meiotic chromosomes of the asexual species *Darevskia dahli*. A) Karyotype with 2n = 38 is widely shared among *Darevskia* lizards. B) Identification of the W sex chromosome by visualization of constitutive heterochromatin which is absent on the pair of microchromosomes (labelled as m). C) Fluorescence *in situ* hybridization (FISH) with the telomeric (TTAGGG)_n_ probe (red) revealed a standard distribution in the terminal regions of all chromosomes. D) The detection of 18S rDNA clusters (green) showed a different amount of accumulation on homeologs. In contrast, the signals on two chromosomes of different sizes, providing evidence of a hybrid origin in the otherwise conserved karyotype of *D. unisexualis* (box). E) All oocytes in the diplotene stage have double the number of chromosomes (i.e., 38 bivalents = 76 chromosomes) compared to the mitotic nuclei, documenting past premeiotic endoreplication. Telomeric FISH was applied to facilitate the counting of chromosomes/bivalents. Note that each detail shows one or more bivalents. The asterisk labels a small bivalent with strong heterochromatic region indicating putative WW bivalent. Scale bar = 10 μm. Complete sets of results for all species studied here are in the Supplementary Material.

The analysis of the diplotene chromosomes in the sexual species *D. r. nairensis* revealed 19 bivalents formed by 38 chromosomes (Supplementary Figure S3). As in mitotic chromosomes, FISH with the telomeric probe did not reveal interstitial telomeric sites and signals were located only in the terminal part of the chromosomes. In all oocytes of the parthenogenetic *D. armeniaca, D. unisexualis* and *D. dahli* we observed 38 bivalents, indicating the presence of 76 chromosomes (Supplementary Figures S4-S6).

In all three obligate parthenogens, we found only oocytes with duplicated genomes in diplotene, suggesting that premeiotic endoreplication is required for progression to later meiotic stages. We can only speculate why the mechanism of the formation of unreduced oocytes was misinterpreted in previous studies of the parthenogenetic lizards of the genus *Darevskia*. Similar to the other parthenogenetic complexes of the hybrid origin studied so far (*Aspidoscelis*: Lutes et al., 2010; Newton et al., 2016; Cole et al., 2020; *Hemiphyllodactylus, Heteronotia* and *Lepidodactylus*: Dedukh et al., 2022), only a small fraction of oocytes undergoes premeiotic duplication and enter later stages also in *Darevskia armeniaca* (Spangenberg et al., 2024). The majority of meiotic oocytes (even well over 90%) have an initial ploidy level, they usually do not pair chromosomes correctly and do not progress through meiosis. It can be confusing that only a small fraction of the cells, easily missed when only a few oocytes are examined, are doing what is essential for reproduction. Along with a pairing of autosomes, Spangenberg et al. (2024) found that the sex chromosomes in around half of the non-duplicated oocytes are not correctly paired at pachytene in the parthenogenetic rock lizards. In contrast, a contact between Z and W sex chromosomes was found in the remaining non-duplicated oocytes, but a support for the synapsis between the homologous chromosomes for example by the visualisation of recombination foci allowing the identification of paired chromosomes during pachytene was not provided. These authors assumed that the sex chromosomes and autosomes are synapsed in these oocytes and that such cells could proceed to the next phases of meiosis. Nevertheless, we found exclusive oocytes with premeiotic endoreplication at the later stages of meiosis (diplotene), although they form only a small fraction at earlier stages. The analysis of diplotene oocytes was earlier performed in the parthenogenetic *D. armeniaca* (Kupriyanova, 2009). However, the obtained chromosomal spreads were either incomplete or bivalents were intensively intermixed thus preventing their counting.

The discovery of the mechanism of oogenesis in the parthenogenetic rock lizards has important implications. Premeiotic endoreplication followed by the pairing and recombination in identical chromosome copies ensures clonality, at least temporarily. In the longer term, a loss of heterozygosity in hybrids with premeiotic endoreplication is most likely due to occasional recombination between orthologs and gene conversion on the homeologs, as documented in gynogenetic teleost fishes (Janko et al., 2021). Gene conversion should occur more likely at recombination hotspots rather than being completely randomly distributed (Lorenz and Mpaulo, 2022). On the other hand, central fusion should be associated with a typical recombination between homologous chromosomes, and thus heterozygosity should be preserved only in the central part of the chromosomes and lost in the regions of the chromosome arms between a recombination site and the telomere of each chromosome. Quite significant loss of heterozygosity was recorded in *Darevskia* in certain microsatellite loci (Tarkhnishvili et al., 2020). It should be tested whether the distribution of homozygous loci throughout the genomes is more consistent with conversion and random occasional recombination between orthologs, or regular recombination of chromosomes expected under the fusion hypothesis by genomic data.

The apparent variability of mechanisms in the obligate parthenogens of the hybrid origin, based on the information available at the time in parthenogenetic *Aspidoscelis* and *Darevskia* lizards, led to the conclusion that “the mechanisms of meiosis in vertebrate parthenogens may not conform to a one-size-fits-all scenario” (Freitas et al., 2022). However, it seems that we need to revise this view: while terminal fusion and postmeiotic doubling connected with the loss of heterozygosity seem to be the mechanism of facultative parthenogenesis (Booth et al., 2012, 2014; Booth and Schuett, 2016; Kratochvíl et al., 2020; Ho et al., 2023), only premeiotic endoreplication is now known in obligate parthenogenetic vertebrates of hybrid origin. The same mechanism was uncovered in the studied representatives of eight or nine independent hybrid origins of obligate parthenogenesis: in the teiid diploids *A. tesselata* (Lutes et al., 2010) (1) and *A. neomexicana* (Newton et al., 2016) (2), in the triploid *A. uniparens* (Cuellar, 1971) and tetraploid *A. priscillae* originating from the laboratory cross of *A. uniparens* with a male of sexual diploid *A. inornatus* (Cole et al., 2017) (3), in the triploid geckos *Heteronotia binoei* (Dedukh et al., 2022) (4) and *Hemiphyllodactylus typus* (Dedukh et al., 2022) (5), in the diploid and triploid geckos *Lepidodactylus lugubris* (Dedukh et al., 2022) (6), and - as shown in this study, in the diploid lacertids *D. unisexualis* (7), *D. dahli* (8), and *D. armeniaca* (9 - depending on the mode of its origin, as it might inherited parthenogenesis from its already parthenogenetic ancestor; Tarkhnishvili et al., 2020).

Our finding of exclusively endoreplicated oocytes in late diplotene, although cells with the original ploidy levels being observed in earlier stages (Spangenberg et al., 2020, 2024), suggests that the pachytene checkpoint is conserved and active in all of the parthenogenetic lizards studied. The conservation of the fully functional pachytene checkpoint may indicate its importance to prevent aneuploidy in both sexual females and parthenogenetic lineages. It seems that parthenogenesis was forced to find a way to produce unreduced eggs while still passing through the strict control of the pachytene checkpoint. It appears that two solutions to this challenge were found during the evolution: premeiotic endoreplication after which all chromosomes are properly paired in hybrids, and normal pairing followed by typical meiosis and a fusion in facultative parthenogenesis.

Previously, we speculated that the repeated involvement of premeiotic endoreplication for the production of unreduced eggs in hybrids might be caused by the co-option of the pre-existing mechanism of induced genome duplication by increased levels of apoptotic cells, specifically by a mass of discarded oocytes with the original ploidy level arrested by the pachytene checkpoint (Dedukh et al., 2022). This hypothesis attempts to explain why premeiotic endoreplication occurs repeatedly and instantaneously in the F1 hybrid generation and, at the same time, why the majority of oocytes enter meiosis without prior endoreplication and fail to complete their development. We also emphasised that premeiotic endoreplication - in contrast to egg/polar body fusion - opens the way to polyploidization: it seems easier to produce unreduced oocytes after incorporation of the other haploid chromosome set using this mechanism than under a fusion. The possible backcrossing leading to the emergence of *D. armeniaca* and other introgressions in the *Darevskia* lizards suggested by Tarkhnishvili et al. (2020) would be more plausible through a fertile triploid produced by a fertilisation of a diploid egg of a parthenogenetic lineage with a haploid sperm of a sexual species under endoreplication than under fusion, as already pointed out by these authors. It should be noted that the observed allele sharing could be the result of a gene flow from the parthenogenetic to the sexual population as known in other asexual complexes through a polyploid: in some teleost hybrid complexes, triploids produce haploid recombined gametes although their diploid ancestries were gynogenetic (Alves et al., 2001; Goddard et al., 1998; Dedukh et al., 2023). However, there is still no direct evidence for fertility of triploid females in *Darevskia* (only for well-developed reproductive organs in them), and no all-triploid parthenogenetic populations have ever been found in this genus. Nevertheless, the finding of tetraploid individuals (Danielyan et al., 2008), probably resulting from a cross between a triploid and a diploid sexual individual, and triploid females showing signs of previous clutch laying in the field suggest that at least some triploid females could be fertile (reviewed in Tarkhnishvili et al., 2020).

In summary, we emphasise that only premeiotic endoreplication has been documented as a mechanism of the production of unreduced eggs in obligate parthenogenetic vertebrates of hybrid origin to date. It seems that evolution has not been so creative in finding ways to escape sexual reproduction in them, always maintaining the pachytene checkpoint and co-opting the same developmental mechanisms, the premeiotic endoreplication, over and over again.

## Acknowledgements

We thank Eugeny Iryshkov and Ilya Brinev for their help in the field and Šárka Pelikánová for continuing support in the laboratory.

## Author contributions

Conceptualization: D.D., M.A., L.K.; methodology: D.D., M.A.; field study: E.G, D.D.; formal analyses: D.D., M.A.; visualisation: D.D., M.A.; resources: D.D., M.A., L.K., R.P., M.A.; data curation: D.D., M.A., L.K.; funding acquisition: D.D., M.A., L.K; writing - original draft: D.D., M.A., L.K.; writing - review & editing: all authors.

## Supplementary material

Table S1, figures S1-S6.

## References

Abdala, C. S., Baldo, D., Juárez, R. A. and Espinoza, R. E. (2016). The first parthenogenetic pleurodont iguanian: A new all-female Liolaemus (Squamata: Liolaemidae) from Western Argentina. Copeia 104(2), 487–497. 10.1643/CH-15-381

Adams, M., Foster, R., Hutchinson, M. N., Hutchinson, R. G. and Donnellan, S. C. (2003). The Australian scincid lizard Menetia greyii: A new instance of widespread vertebrate parthenogenesis. Evolution 57(11), 2619–2627. 10.1111/j.0014-3820.2003.tb01504.x

Altmanová, M., Doležálková-Kaštánková, M., Jablonski, D., Strachinis, I., Vergilov, V., Vacheva, E., Iannucci, A., Choleva, L., Ráb, P., Moravec, J. et al. (2023). Karyotype stasis but species-specific repetitive DNA patterns in Anguis lizards (Squamata: Anguidae), in the evolutionary framework of Anguiformes. Zool. J. Linn. Soc., zlad153. 10.1093/zoolinnean/zlad153

Alves, M. J., Coelho, M. M. and Collares-Pereira, M. J. (2001). Evolution in action through hybridisation and polyploidy in an Iberian freshwater fish: A genetic review. Genetica 111(1-3), 375–385. 10.1023/a:1013783029921

Arakelyan, M., Spangenberg, V., Petrosyan, V., Ryskov, A., Kolomiets, O. and Galoyan, E. (2023). Evolution of parthenogenetic reproduction in Caucasian rock lizards: A review. Curr. Zool. 69(2), 128–135. 10.1093/cz/zoac036

Avise, J. C. (2008). Clonality: The Genetics, Ecology, and Evolution of Sexual Abstinence in Vertebrate Animals. New York: Oxford Academic (online edn, 1 Jan. 2009). 10.1093/acprof:oso/9780195369670.001.0001

Barley, A. J., Reeder, T. W., Nieto-Montes De Oca, A., Cole, C. J. and Thomson, R. C. (2021). A new diploid parthenogenetic whiptail lizard from Sonora, Mexico, is the ‘missing link’ in the evolutionary transition to polyploidy. Am. Nat. 198(2), 295–309. 10.1086/715056

Bogart, J. P., Bartoszek, J., Noble, D. W. A. and Bi, K. (2009). Sex in unisexual salamanders: Discovery of a new sperm donor with ancient affinities. Heredity 103, 483–493. 10.1038/hdy.2009.83

Booth, W. and Schuett, G. L. (2016). The emerging phylogenetic pattern of parthenogenesis in snakes. Biol. J. Linn. Soc. 118(2), 172–186. 10.1111/bij.12744

Booth, W., Schuett, G. W., Ridgway, A., Buxton, D. W., Castoe, T. A., Bastone, G., Bennett, C. and McMahan, W. (2014). New insights on facultative parthenogenesis in pythons. Biol. J. Linn. Soc. 112(3), 461–468. 10.1111/bij.12286

Booth, W., Smith, C. F., Eskridge, P. H., Hoss, S. K., Mendelson, J. R. and Schuett, G. W. (2012). Facultative parthenogenesis discovered in wild vertebrates. Biol. Lett. 8(6), 983–985. 10.1098/rsbl.2012.0666

Brunes, T. O., da Silva, A. J., Marques-Souza, S., Rodrigues, M. T. and Pellegrino, K. C. M. (2019). Not always young: The first vertebrate ancient origin of true parthenogenesis found in an Amazon leaf litter lizard with evidence of mitochondrial haplotypes surfing on the wave of a range expansion. Mol. Phylogenet. Evol. 135, 105–122. 10.1016/j.ympev.2019.01.023

Gall, J. G., Murphy, C., Callan, H. G. and Wu, Z. A. (1991). Lampbrush chromosomes. Methods Cell Biol. 36, 149–166.

Cole, C. J., Dessauer, H. C., Paulissen, M. A. and Walker, J. M. (2020). Hybridization between whiptail lizards in Texas: Aspidoscelis laredoensis and A. gularis, with notes on reproduction of a hybrid. Am. Mus. Novit. 2020, 1–13. 10.1206/3947.1

Cole, C. J., Taylor, H. L., Neaves, W. B, Baumann, D. P., Newton, A., Schnittker, R. and Baumann, P. (2017). The second known tetraploid species of parthenogenetic tetrapod (Reptilia: Squamata: Teiidae): Description, reproduction, comparisons with ancestral taxa, and origins of multiple clones. Bull. Mus. Comp. Zool. 161(8), 285–321. 10.3099/MCZ37.1

Cuellar, O. (1971) Reproduction and the mechanism of meiotic restitution in the parthenogenetic lizard Cnemidophorus uniparens. J. Morphol. 133(2), 139–165. 10.1002/jmor.1051330203

Danielyan, F., Arakelyan, M. and Stepanyan, I. (2008). Hybrids of Darevskia valentini, D. armeniaca and D. unisexualis from a sympatric population in Armenia. Amphibia-Reptilia 29(4), 487–504. 10.1163/156853808786230424

Darevsky, I. S. (1966). Natural parthenogenesis in a polymorphic group of Caucasian rock lizards related to Lacerta saxicola Eversmann. Journal of the Ohio Herpetological Society 5(4), 115–152.

Darevsky, I. S., Kupriyanova, L. A. and Uzzell, T. (1985). Parthenogenesis in reptiles. In Biology of the Reptilia, Vol. 15 (eds. C. Gans and F. Billet), pp. 412–526. New York: John Wiley and Sons.

Dedukh, D., Altmanová, M., Klíma, J. and Kratochvíl, L. (2022). Premeiotic endoreplication is essential for obligate parthenogenesis in geckos. Development 149(7), dev200345. 10.1242/dev.200345

Dedukh, D., Marta, A., Myung, R.-Y., Ko, M.-H., Choi, D.-S., Won, Y.-J. and Janko, K. (2023). From asexuality to sexual reproduction: Cyclical switch of gametogenic pathways in hybrids depends on ploidy level. BioRxiv 2023.06.18.545483. 10.1101/2023.06.18.545483

Freitas, S. N., Harris, D. J., Sillero, N., Arakelyan, M., Butlin, R. K. and Carretero, M. A. (2019). The role of hybridisation in the origin and evolutionary persistence of vertebrate parthenogens: A case study of Darevskia lizards. Heredity (Edinb.) 123(6), 795–808. 10.1038/s41437-019-0256-5

Freitas, S., Westram, A. M., Schwander, T., Arakelyan, M., Ilgaz, Ç., Kumlutas, Y., Harris, D. J., Carretero, M. A. and Butlin, R. K. (2022). Parthenogenesis in Darevskia lizards: A rare outcome of common hybridization, not a common outcome of rare hybridization. Evolution 76(5), 899–914. 10.1111/evo.14462

Goddard, K. A., Megwinoff, O., Wessner, L. L. and Giaimo, F. (1998) Confirmation of gynogenesis in Phoxinus eos-neogaeus (Pisces: Cyprinidae). J. Hered. 89(2), 151–157. 10.1093/jhered/89.2.151

Grismer, J. L., Bauer, A. M., Grismer, L. L., Thirakhupt, K., Aowphol, A., Oaks, J. R., Wood, P. L., Onn, C. K., Thy, N., Cota, M. et al. (2014). Multiple origins of parthenogenesis, and a revised species phylogeny for the Southeast Asian butterfly lizards, Leiolepis. Biol. J. Linn. Soc. 113(4), 1080–1093. 10.1111/bij.12367

Ho, D. V., Tormey, D., Odell, A., Newton, A. A., Schnittker, R. R., Baumann, D. P., Neaves, W. B., Schroeder, M. R., Sigauke, R. F., Barley, A. J. et al. (2023). Post-meiotic mechanism of facultative parthenogenesis in gonochoristic whiptail lizard species. BioRxiv 2023.09.21.558237. 10.1101/2023.09.21.558237

Janko, K., Bartoš, O., Kočí, J., Roslein, J., Drdová, E. J., Kotusz, J., Eisner, J., Mokrejš, M. and Štefková-Kašparová, E. (2021). Genome fractionation and loss of heterozygosity in hybrids and polyploids: Mechanisms, consequences for selection, and link to gene function. Mol. Biol. Evol. 38(12), 5255–5274. 10.1093/molbev/msab249

Kratochvíl, L., Vukić, J., červenka, J., Kubička, L., Johnson Pokorná, M., Kukačková, D., Rovatsos, M. and Piálek, L. (2020). Mixed-sex offspring produced via cryptic parthenogenesis in a lizard. Mol. Ecol. 29(21), 4118–4127. 10.1111/mec.15617

Kupriyanova, L. (2009). Cytogenetic and genetic trends in the evolution of unisexual lizards. Cytogenet. Genome Res. 127(2-4), 273–279. 10.1159/000303325

Kupriyanova, L. A., Safronova, L. D., Sycheva, V. B., Danielyan, F. D. and Petrosyan, V. G. (2021). Oogenesis (prophase 1 of meiosis) and mitotic chromosomes of parthenogenetic species Darevskia armeniaca (family Lacertidae). Biol. Bull. 48, 274–280. 10.1134/S1062359021030080

Lamatsch, D. K. and Stöck, M. (2009). Sperm-dependent parthenogenesis and hybridogenesis in teleost fishes. In Lost Sex, pp. 399-432. Dordrecht: Springer Netherlands.

Lavanchy, G. and Schwander, T. (2019). Hybridogenesis. Curr. Biol. 29(1), R9–R11. 10.1016/j.cub.2018.11.046

Lorenz, A. and Mpaulo, S. J. (2022). Gene conversion: A non-Mendelian process integral to meiotic recombination. Heredity 129(1), 56–63. 10.1038/s41437-022-00523-3

Lutes, A. A., Neaves, W. B., Baumann, D. P., Wiegraebe, W. and Baumann, P. (2010). Sister chromosome pairing maintains heterozygosity in parthenogenetic lizards. Nature 464, 283–286. 10.1038/nature08818

Murphy, R. W., Fu, J., MacCulloch, R. D., Darevsky, I. S. and Kuprianova, L. A. (2000). A fine line between sex and unisexuality: the phylogenetic constraints on parthenogenesis in lacertid lizards. Zool. J. Linn. Soc. 130(4), 527–549. 10.1111/j.1096-3642.2000.tb02200.x

Newton, A. A., Schnittker, R. R., Yu, Z., Munday, S. S., Baumann, D. P., Neaves, W. B. and Baumann, P. (2016). Widespread failure to complete meiosis does not impair fecundity in parthenogenetic whiptail lizards. Development 143(23), 4486–4494. 10.1242/dev.141283

Poignet, M., Johnson Pokorná, M., Altmanová, M., Majtánová, Z., Dedukh, D., Albrecht, T., Reif, J., Osiejuk, T. and Reifová, R. (2021). Comparison of karyotypes in two hybridizing passerine species: Conserved chromosomal structure but divergence in centromeric repeats. Front. Genet. 12, 768987. 10.3389/fgene.2021.768987

Reeder, T. W., Cole, C. J. and Dessauer, H. C. (2002). Phylogenetic relationships of whiptail lizards of the genus Cnemidophorus (Squamata: Teiidae): A test of monophyly, reevaluation of karyotypic evolution, and review of hybrid origins. Am. Mus. Novit. 2002(3365), 1–61. 10.1206/0003-0082(2002)365<0001:PROWLO>2.0.CO;2

Sinclair, E. A., Pramuk, J. B., Bezy, R. L., Crandall, K. A. and Sites, J. W. Jr (2010). DNA evidence for nonhybrid origins of parthenogenesis in natural populations of vertebrates. Evolution 64(5), 1346–1357. 10.1111/j.1558-5646.2009.00893.x

Spangenberg, V., Arakelyan, M., Cioffi, M. B., Liehr, T., Al-Rikabi, A., Martynova, E., Danielyan, F., Stepanyan, I., Galoyan, E. and Kolomiets, O. (2020). Cytogenetic mechanisms of unisexuality in rock lizards. Sci. Rep. 10, 8697. 10.1038/s41598-020-65686-7

Spangenberg, V., Arakelyan, M., Galoyan, E., Martirosyan, I., Bogomazova, A., Martynova, E., Cioffi, M. B., Liehr, T., Al-Rikabi, A, Osipov, F. et al. (2021). Meiotic synapsis of homeologous chromosomes and mismatch repair protein detection in the parthenogenetic rock lizard Darevskia unisexualis. Mol. Reprod. Dev. 88(2), 119–127. 10.1002/mrd.23450

Spangenberg, V., Arakelyan, M., Simanovsky, S., Khachatryan, E. and Kolomiets, O. (2024). Tendency towards clonality: Deviations of meiosis in parthenogenetic Caucasian rock lizards. PREPRINT (Version 2) available at Research Square. 10.21203/rs.3.rs-3936576/v2

Stöck, M., Dedukh, D., Reifová, R., Lamatsch, D. K., Starostová, Z. and Janko, K. (2021a). Sex chromosomes in meiotic, hemiclonal, clonal and polyploid hybrid vertebrates: Along the ‘extended speciation continuum’. Philos. Trans. R. Soc. B 376(1833), 20200103. 10.1098/rstb.2020.0103

Stöck, M., Kratochvíl, L., Kuhl, H., Rovatsos, M., Evans, B. J., Suh, A., Valenzuela, N., Veyrunes, F., Zhou, Q., Gamble, T. et al. (2021b). A brief review of vertebrate sex evolution with a pledge for integrative research: towards ‘sexomics’. Phil. Trans. R. Soc. B 376(1832), 20200426. 10.1098/rstb.2020.0426

Tarkhnishvili, D., Yanchukov, A., Şahin, M. K., Gabelaia, M., Murtskhvaladze, M., Candan, K., Galoyan, E., Arakelyan, M., Iankoshvili, G., Kumlutaş, Y. et al. (2020). Genotypic similarities among the parthenogenetic Darevskia rock lizards with different hybrid origins. BMC Evol. Biol. 20, 122. 10.1186/s12862-020-01690-9

Uzzell, T. (1970). Meiotic mechanisms of naturally occurring unisexual vertebrates. Am. Nat. 104(939), 433–445.

Vergun, A. A., Girnyk, A. E., Korchagin, V. I., Semyenova, S. K., Arakelyan, M. S., Danielyan, F. D., Murphy, R. W. and Ryskov A. P. (2020). Origin, clonal diversity, and evolution of the parthenogenetic lizard Darevskia unisexualis. BMC Genom. 21, 351. 10.1186/s12864-020-6759-x

Wynn, A. H., Cole, C. J. and Gardner, A. L. (1987). Apparent triploidy in the unisexual brahminy blind snake, Ramphotyphlops braminus. Am. Mus. Novit. 2868, 1–7.

Yanchukov, A., Tarkhnishvili, D., Erdolu, M., Şahin, M. K., Candan, K., Murtskhvaladze, M., Gabelaia, M., Iankoshvili, G., Barateli, N., Ilgaz, C. et al. (2022). Precise paternal ancestry of hybrid unisexual ZW lizards (genus Darevskia: Lacertidae: Squamata) revealed by Z-linked genomic markers. Biol. J. Linn. Soc. 136(2), 293–305. 10.1093/biolinnean/blac023

