## Supplementary material for "Premeiotic endoreplication is the mechanism of obligate parthenogenesis in rock lizards of the genus *Darevskia*"

**Table S1.** *Darevskia* species analysed in this study, their reproduction mode, origin of individuals and methods used. The parental (maternal × paternal) species of unisexual lineages according to Yanchukov et al. (2022). F – female. FISH – fluorescence *in situ* hybridization.

| **Species** | **Reproduction mode** | **Individual** | **Locality (Armenia)** | **Coordinates** | **Number of analysed diplotene oocytes** | **Mitotic karyotyping,  C-banding, 18S rDNA FISH, telomeric FISH** |
| --- | --- | --- | --- | --- | --- | --- |
| *D. raddei nairensis* | Sexual | F1 | Hrazdan | 40.50615N, 44.74847E | 3 | ✓ |
|  |  | F2 | Vagramberd | 40.84278N, 43.75583E | 3 | ✓ |
| *D. armeniaca* | Parthenogenetic | F1 | Dilijan | 40.69720N, 44.85640E | 5 | ✓ |
| (*D. mixta* × *D. valentini*) |  | F2 | Dilijan | 40.69720N, 44.85640E | 3 | ✓ |
| *D. dahli* | Parthenogenetic | F1 | Agvi | 41.07435N, 44.58764E | 2 | ✓ |
| (*D. mixta* × *D. portschinskii*) |  | F2 | Agvi | 41.07435N, 44.58764E | 3 | ✓ |
| *D. unisexualis* | Parthenogenetic | F1 | Hrazdan | 40.50615N, 44.74847E | 2 | – |
| (*D. r. nairensis* × *D. valentini*) |  | F2 | Hrazdan | 40.50615N, 44.74847E | 2 | ✓ |

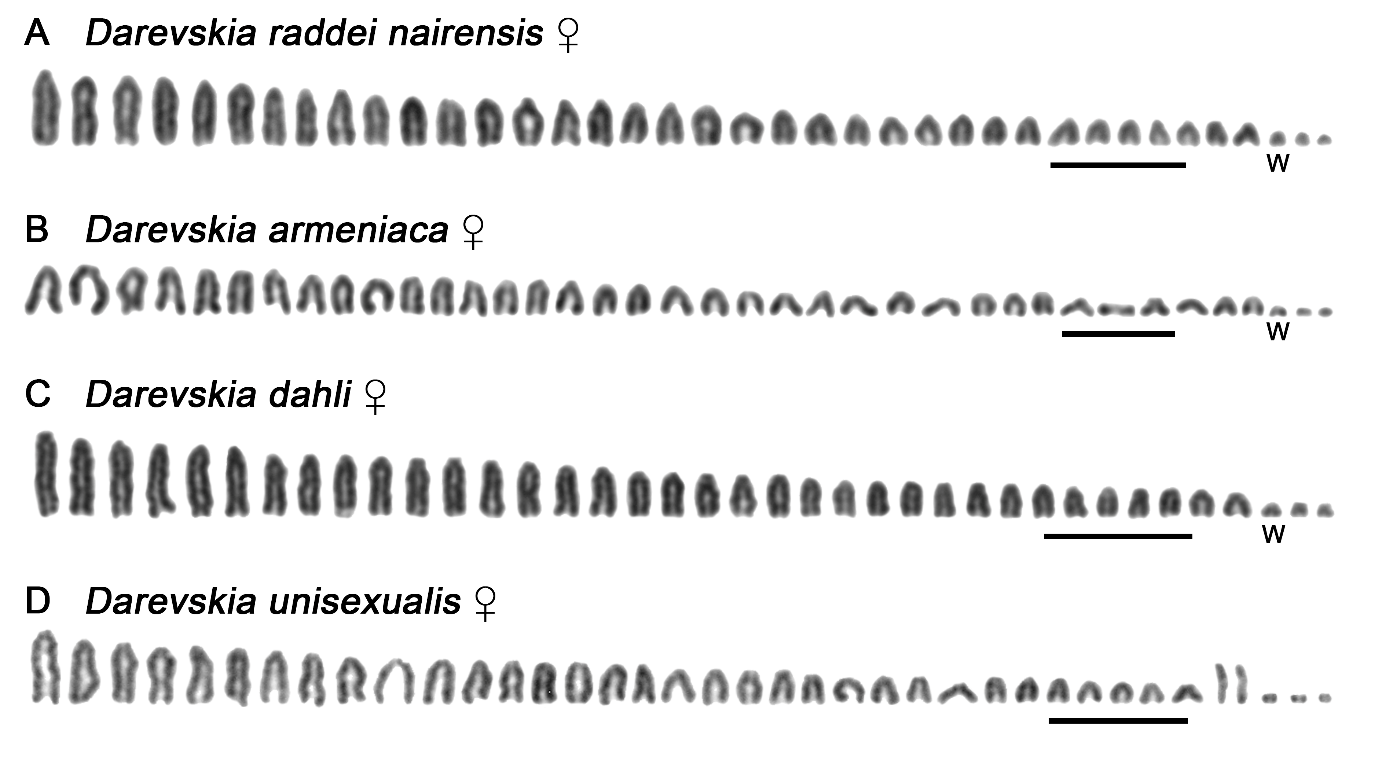

**Figure S1.** Female karyograms of the analysed *Darevskia* species. All tested individuals share the karyotype with 2n = 38. Where identifiable, the sex chromosome W is indicated (its identity was confirmed by the results of sequential C-banding). Scale bar = 10 µm.

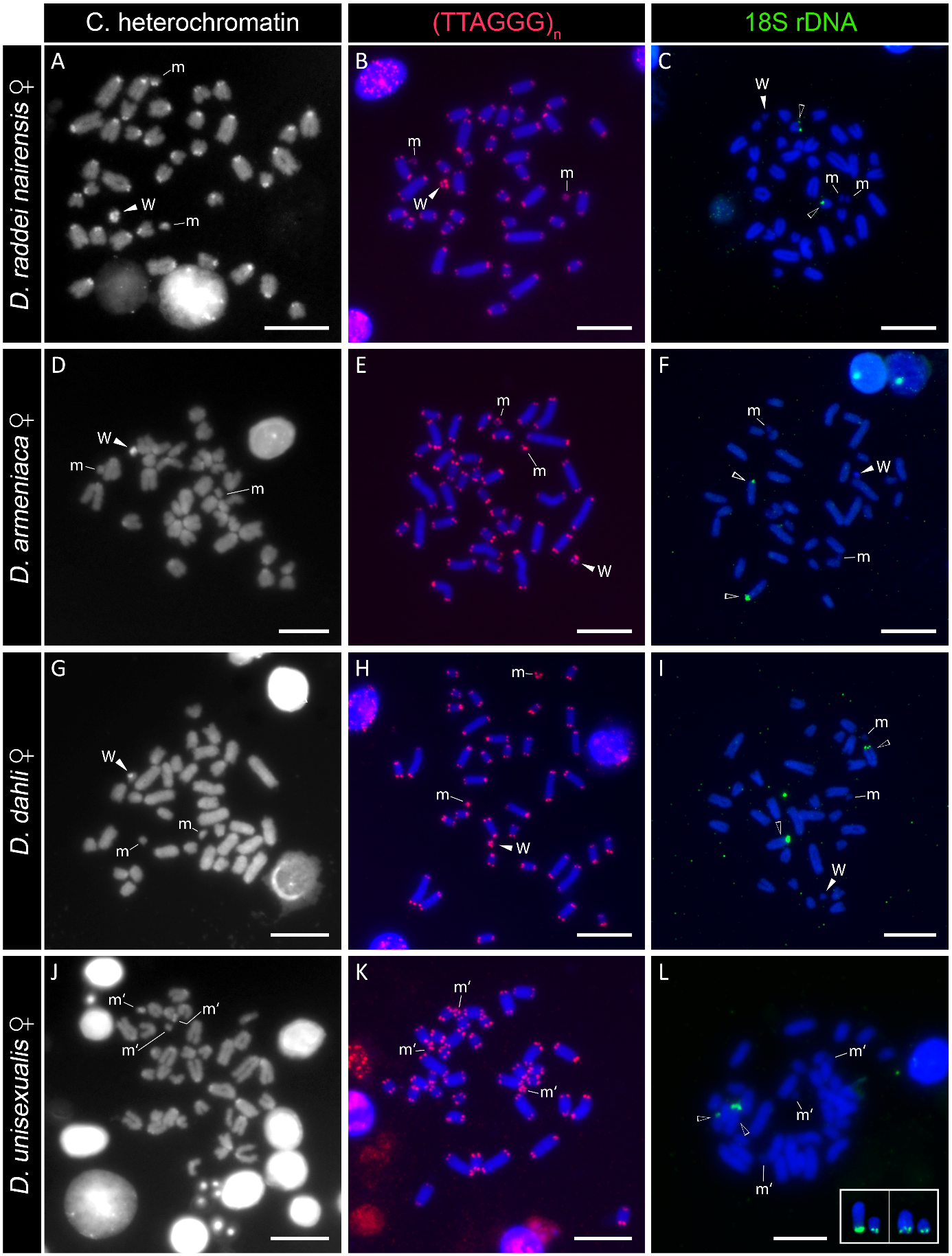

**Figure S2.** Distribution of constitutive heterochromatin (A, D, G, J), telomeric sequences (red; B, E, H, K), and 18S rDNA clusters (green, empty arrowheads; C, F, I, L) on mitotic metaphase chromosomes in *Darevskia* females. Where identifiable, the sex chromosome W is indicated, m assigns an autosomal pair of microchromosomes. In *D. unisexualis* (J–L), m’ labels all three microchromosomes, one of them corresponds to the W chromosome. Note that in *D. unisexualis* the hybridization pattern documents different topology of 18S rDNA clusters between parental species sets of chromosomes, visualized also on chromosomes from other nuclei (box; L). Scale bar = 10 µm.

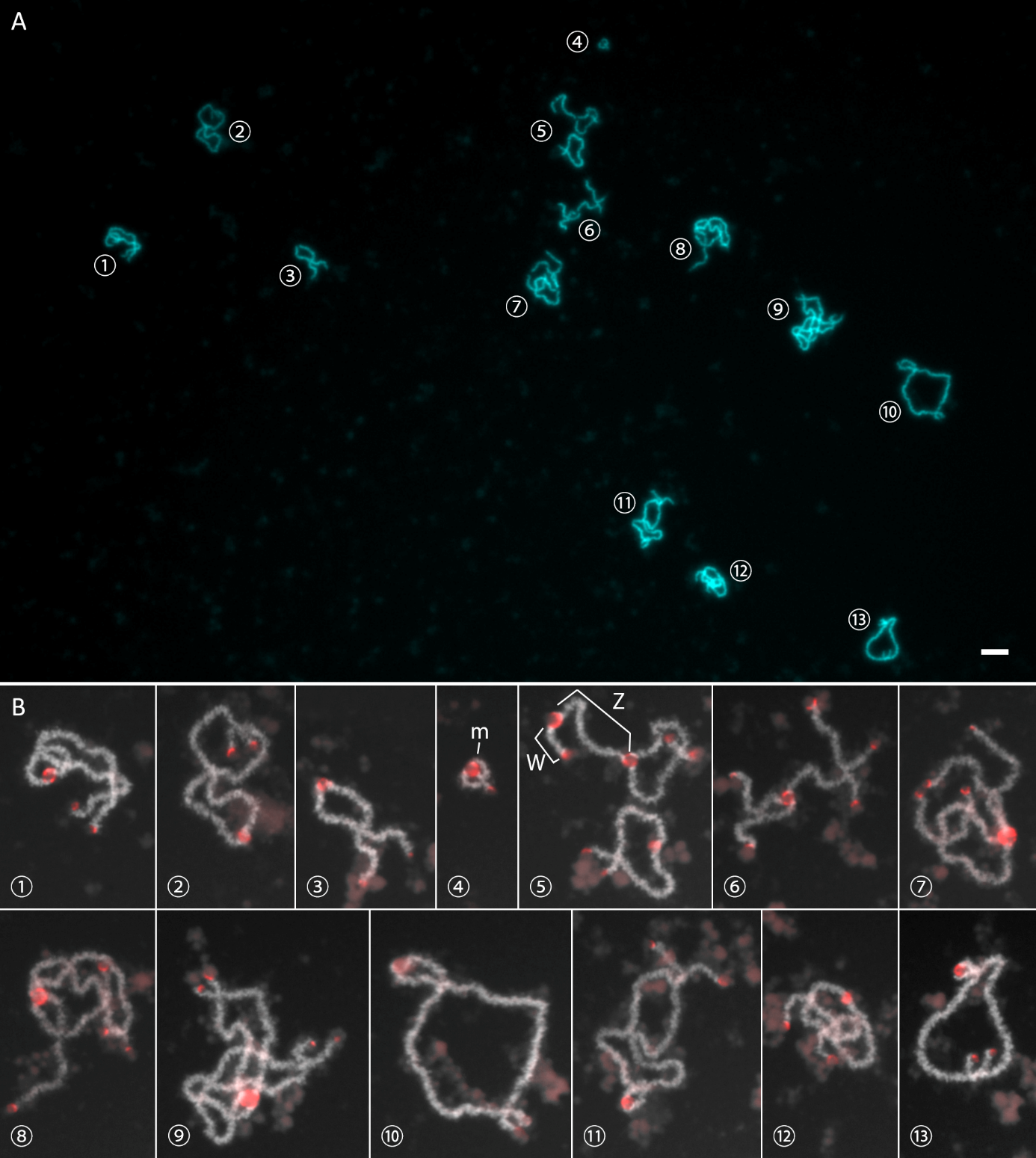

**Figure S3.** Diplotene chromosomes forming 19 bivalents (e.i.38 chromosomes) in the sexual species *D. raddei nairensis* stained by DAPI (A) and with detected telomeres (red, B). Corresponding bivalents in A and B are labelled with the same numbers. In B, m labels bivalent of microchromosomes and the detail no. 5 displays Z and W sex chromosomes and other two bivalents. Whereas all autosomes show chiasmata, the Z and W sex chromosomes are associated only in the telomeric region. Scale bar = 10 µm.

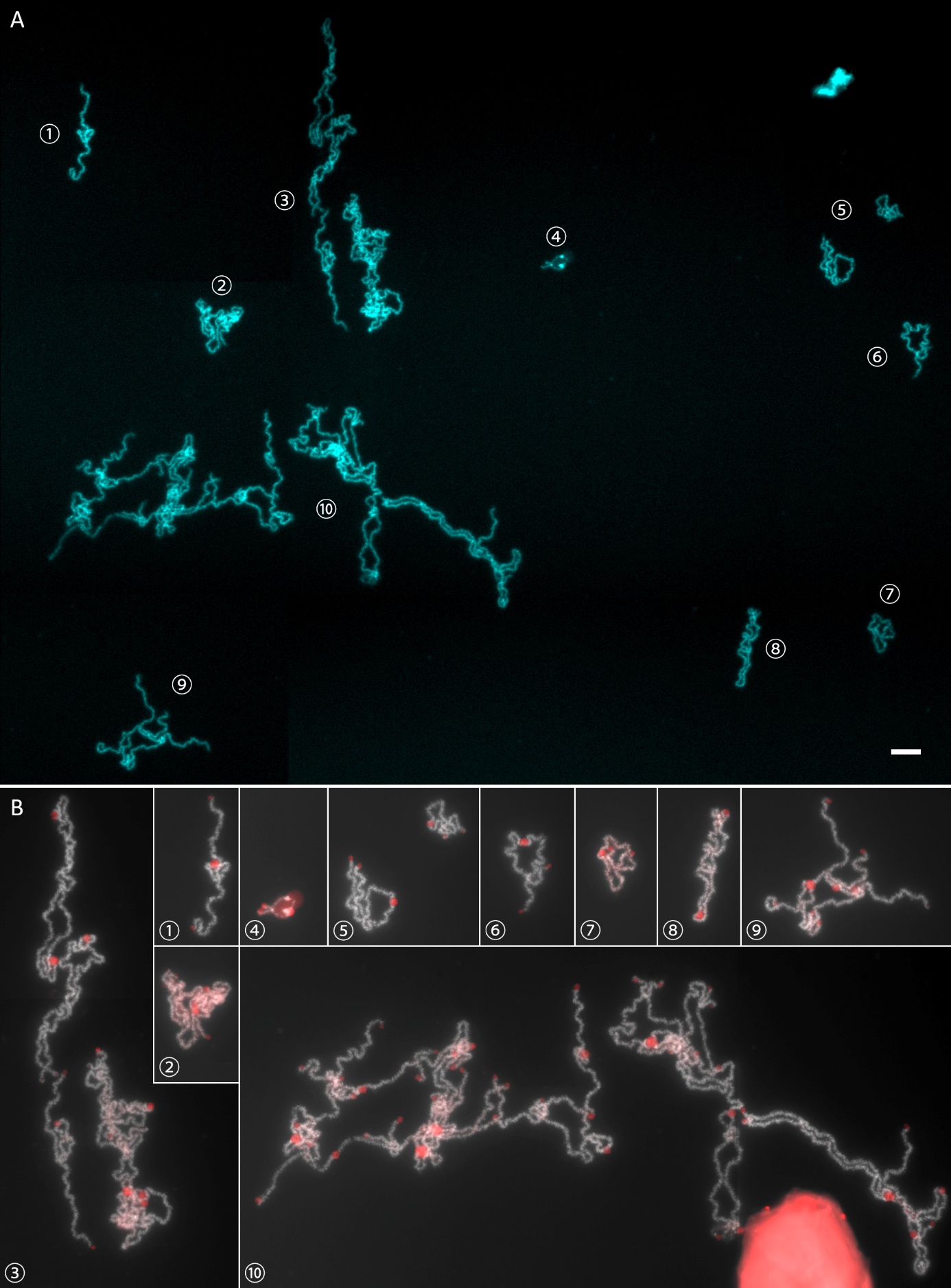

**Figure S4.** Diplotene chromosomes of parthenogenetic species *D. armeniaca* stained by DAPI (A) and with detected telomeres (red, B). The numbers label corresponding bivalent(s) in both A and B. Note the double number of chromosomes (i.e., 38 bivalents = 76 chromosomes) compared to sexual species. Due to many overlaps, it was not possible to distinguish bivalents of W chromosomes and microchromosomes, however one of those bivalents (no. 4, putative WW bivalent) is shown in detail. Scale bar = 10 µm.

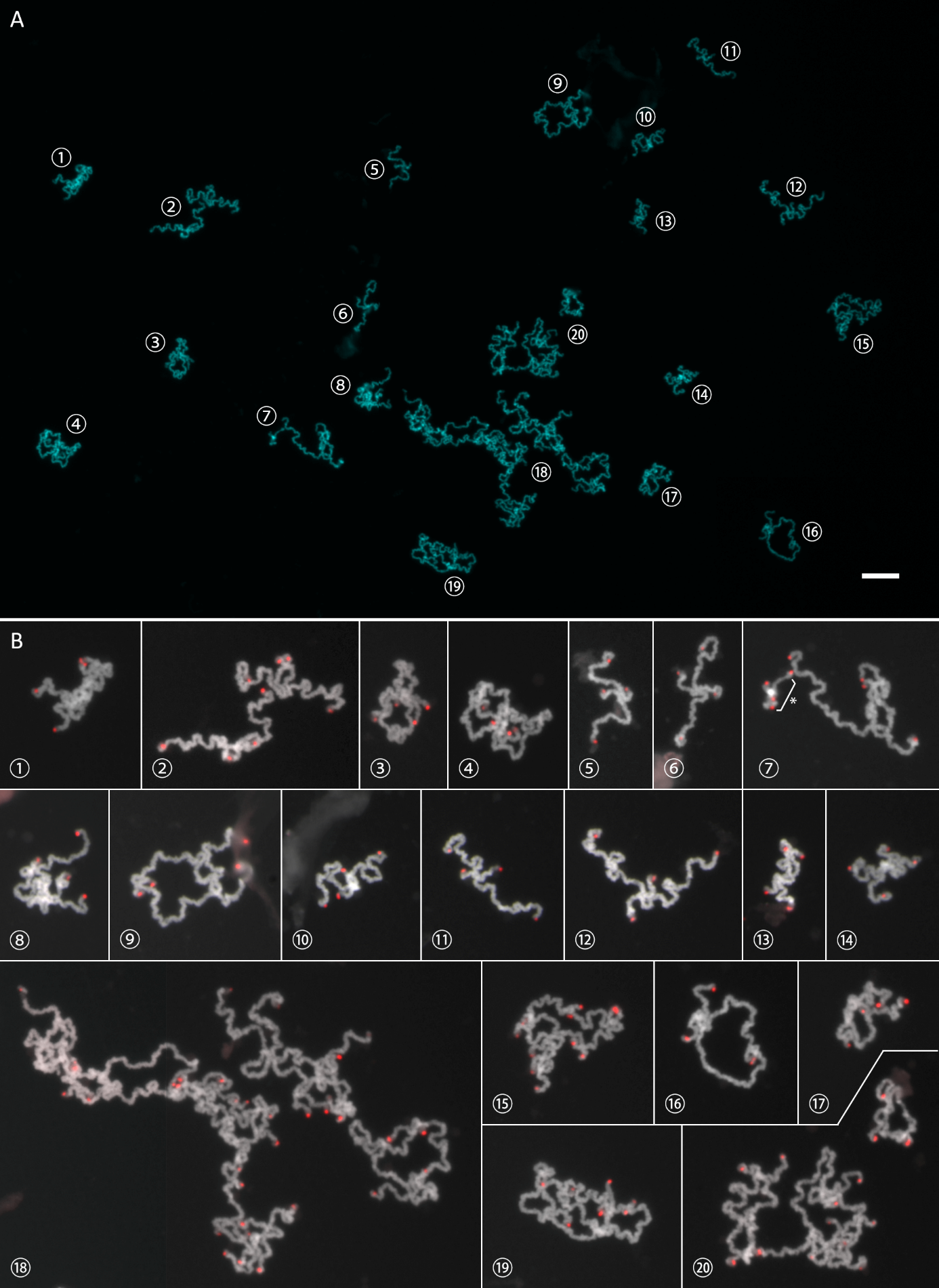
**Figure S5.** Diplotene chromosomes of parthenogenetic species *D. dahli* stained by DAPI (A) and with detected telomeres (red, B). The numbers label corresponding bivalent(s) between A and B. Note the double number of chromosomes (i.e., 38 bivalents = 76 chromosomes) compared to sexual species. Due to many overlaps, it was not possible to distinguish all microchromosomal bivalents; however, one of those bivalents (detail no. 7, marked with an asterisk) had strong heterochromatic region indicating putative WW bivalent. Scale bar = 10 µm.

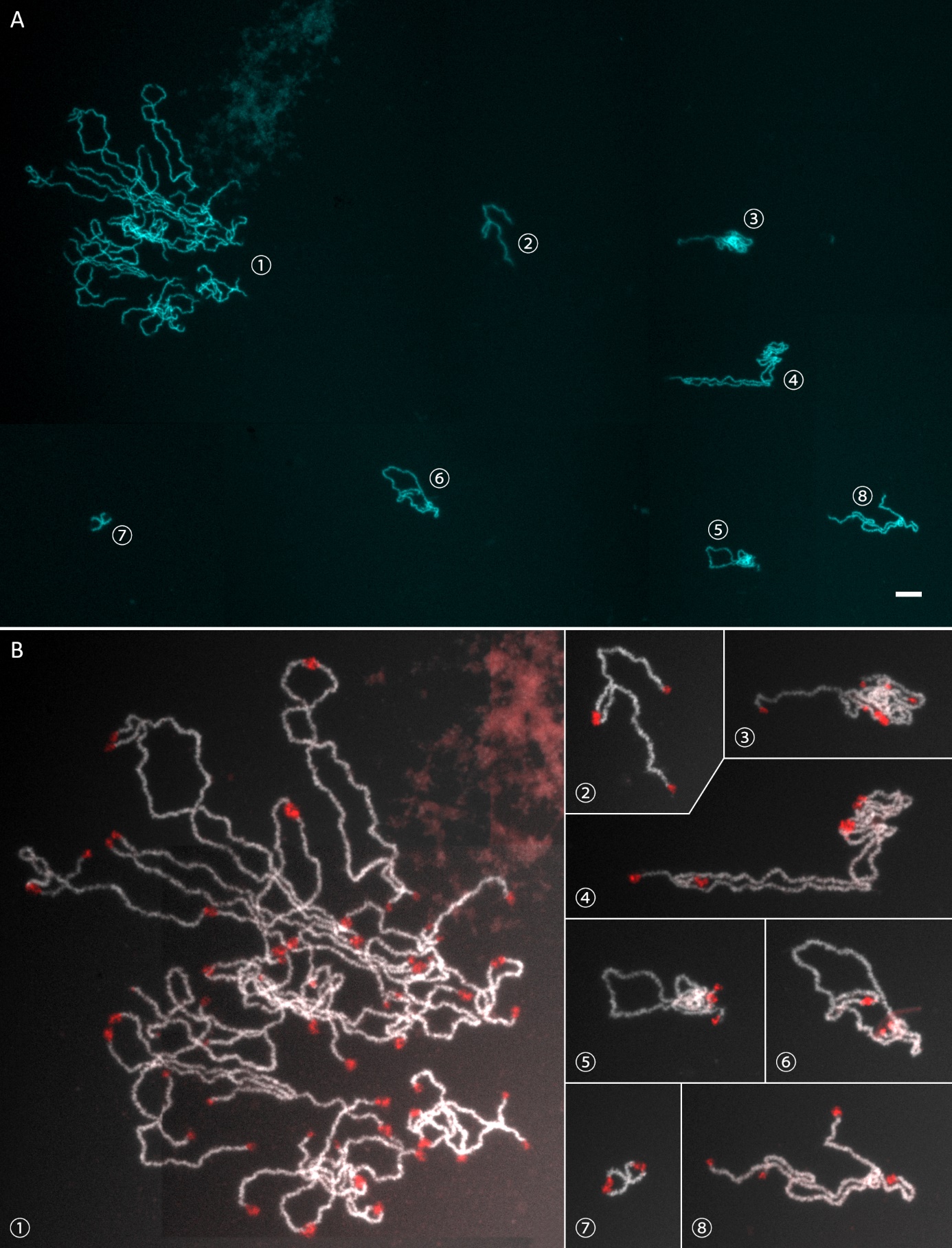
**Figure S6.** Diplotene chromosomes of parthenogenetic species *D. unisexualis* stained by DAPI (A) and with detected telomeres (red, B). The numbers label corresponding bivalent(s) in A and B. Note the double number of chromosomes (i.e., 38 bivalents = 76 chromosomes) compared to sexual species. Due to many overlaps, it was not possible to distinguish bivalents of W chromosomes and microchromosomes; however, one of those bivalents (no. 7) is shown in detail. Scale bar = 10 µm.
